# Fermented foods restructure gut microbiota and promote immune regulation via microbial metabolites

**DOI:** 10.1101/2022.05.11.490523

**Authors:** Sean Paul Spencer, Evelyn Giselle Lemus Silva, Elisa Benedetti Caffery, Matthew Merrill Carter, Rebecca Neal Culver, Min Wang, Rebecca Hope Gellman, Hannah Constance Wastyk, Steven Kyle Higginbottom, Justin Laine Sonnenburg

## Abstract

Fermented foods are ancient and ubiquitous, thought to be consumed in nearly every culture over the last 10,000 years and as part of the hominin diet for millions of years. A growing body of evidence supports their potential health benefits, but the mechanistic basis of their effects on the gut microbiome and host immunity remain to be elucidated. Fermented foods are diverse, each representing a complex mixture of food, microbes, and metabolites creating a significant challenge to disentangle the effects of individual components. Herein, we further define the chemical signature of individual fermented foods to categorize them based on the primary metabolic end-products of fermentation. Using mouse models, we find that fermented foods have both microbiome directed as well as differential host directed effects that correspond to their metabolite composition. Fermented food brine drink shows site-specific restructuring of the gut microbiome and promotion of tolerogenic barrier immunity; fractionation of the brine to examine the effects of the microbe-free, metabolite rich supernatant shows similar activity. Lactate, the main metabolite of lactic acid fermentation and the major metabolite within the brine drink, when administered in water, fuels a trans-kingdom metabolic network to selectively promote the growth of *Akkermansia muciniphila*. in the small intestine, while promoting immune tolerance via an increase in microbiota-dependent Regulatory T-cells. These findings suggest that the beneficial effects of fermented food consumption can be mediated by microbial metabolites within fermented foods, independent of microbial content, and highlight the importance of further defining the diverse chemical landscape of fermented foods to inform their potential health benefits and therapeutic use.

## Introduction

Fermented foods are widely consumed and, prior to industrialization, were a staple of diets around the world^1^. Humans have a deep evolutionary relationship and adaptations to fermented dietary components evidenced by positive selection of alcohol dehydrogenase in fruit-foraging great ape ancestors dating to ~10M years ago, suggesting fermented food consumption by hominins long before human directed fermentation^2^. The current definition of fermented foods is: “foods made through desired microbial growth and enzymatic conversions of food components”^3^. The health benefits of fermented foods remain largely untested in clinical trials^4, 5^, however, recent human studies have highlighted the potential benefits of fermented foods to reduce inflammation and promote microbiome diversity^6, 7^. Further, mechanistic insights into the effect of fermented foods on host biology have been limited by their microbial diversity and chemical complexity^8–14^. The chemical identities of the diverse microbial metabolites within fermented foods are less well described and may be key in understanding their health effects.

The intestinal immune system is exquisitely tuned to respond to microbial and metabolic cues within the GI tract and prior data has demonstrated fermented food administration can have immune impacts^15^. We hypothesized that fermented foods may affect host immunity both via the live microbial component as well as their chemical metabolites. We administered whole fermented foods to mice and directly compared their effects to sterilized fermented foods that contained chemical metabolites in the absence of live organisms.

We used a recently developed microbiome metabolite focused LC-MS pipeline to survey the chemical diversity of common fermented foods^16^ in combination with GC-MS assessment of SCFA and organic acid content. We find that fermented foods can be grouped based on their detailed metabolite profiles, which also corresponds to classification based on their dominant metabolites: lactate and/or acetate. Both lactate and acetate are known to interact with the immune system, however, their role in the context of orally consumed fermented foods has not been well studied. Herein, we characterize the microbial and chemical composition of common fermented foods and assess their impact on microbiota composition and host immunity using mouse models. We find lactate is a keystone metabolite that dominantly restructures intestinal microbiota composition and alters host immunity.

## Results

To investigate the chemical composition of common fermented foods we purchased commercially available fermented foods and assessed the levels of carboxylic acids, including short chain fatty acids (SCFA) (Fig 1a). Lactate was present in high amounts in yogurt and fermented sauerkraut brine, while absent from kombucha and miso. The only SCFA present in significant amounts was acetate, which was found in fermented sauerkraut brine and was the sole carboxylic acid present in kombucha. Miso paste did not have appreciable levels of common carboxylic acid-containing products of fermentation. We employed a wider metabolite survey using a recently developed microbiome-focused liquid chromatography–mass spectrometry (LC-MS) pipeline to identify microbial metabolites present in fermented foods^16^. We found that fermented foods have differing chemical signatures that cluster by type of food fermented and by type of fermentation (acetogenic vs lactic acid) (Figure 1b).

**Figure 1:**
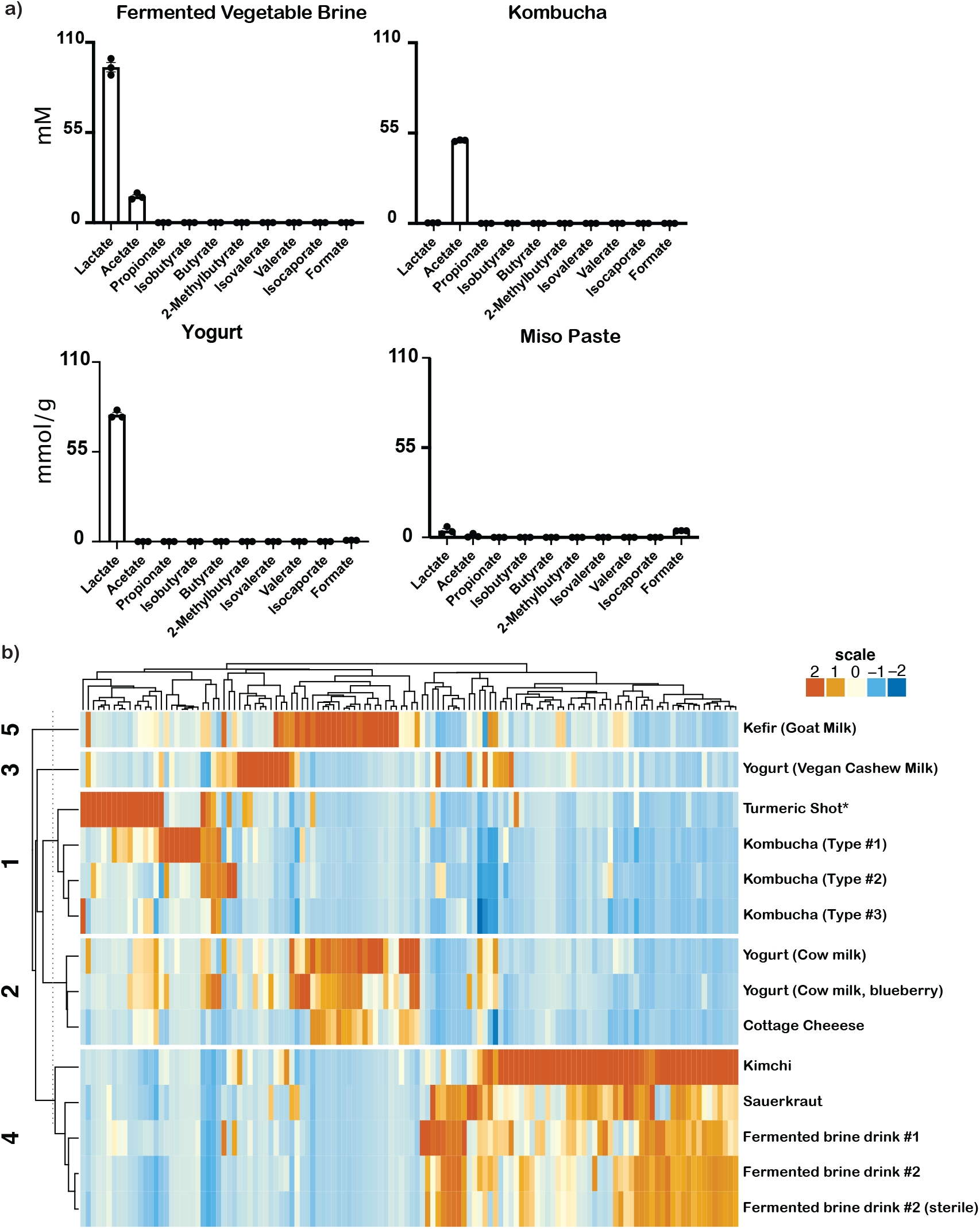
Fermented foods have distinct chemical signatures. a) Organic acid concentrations of commercially available fermented foods (*indicates food was not fermented). b) Hierarchical clustering of metabolites detected across commercially fermented foods. The metabolites with relative abundance in the top 5% of all metabolites found in at least 1 fermented foods are displayed.

As our previous study indicated that fermented foods may alter intestinal microbiota composition^6^, we treated mice with a conventional microbiota with either whole sauerkraut brine, filter sterilized sauerkraut brine, or lactate (Figure 2a). The fecal and cecal bacterial communities of each treatment ordinate separately by permutational multivariate analysis of variance (PERMANOVA), demonstrating that bacterial metabolites as well as lactate alone significantly alter intestinal microbiota composition distinctly from whole sauerkraut brine (Figure 2a).

**Figure 2:**
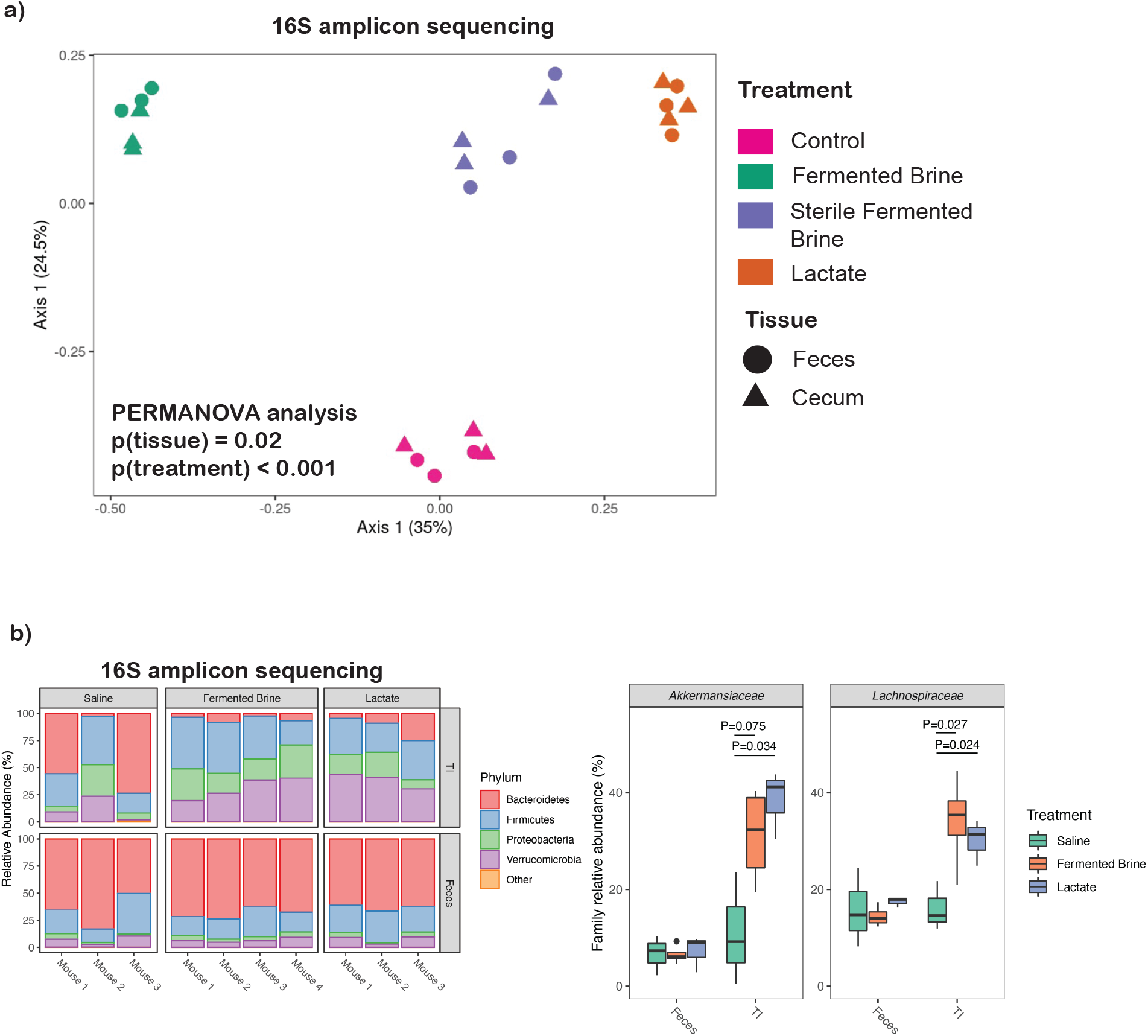
Components of fermented food differentially alter the intestinal microbiome. a) Conventional mice given indicated treatment for 5 weeks and analyzed by 16S amplicon sequencing of feces and cecal contents. Graph displayed is a Principal Coordinate Analysis (PCoA) analyzed via PERMANOVA. b) Gnotobiotic mice colonized with healthy human stool given the indicated treatment for 4 weeks and analyzed via 16S amplicon sequencing with display of relative abundance at the phylum level (left panel) and significantly altered families of bacteria (right panel).

We then assessed the impact of whole sauerkraut brine and lactate on the intestinal microbiota composition using gnotobiotic mice colonized with a healthy human fecal sample (Figure 2b). While the fecal bacterial composition was not significantly altered, the ileal composition was, with enrichments in the family *Akkermansiaceae* and *Lachnospiraceae* (Figure 2b). These data suggest a dominant effect of fermented food intake on the microbiota composition of the small intestine.

Given our data suggesting that fermented food intake alters the small intestinal microbiota, we isolated immune cells from the large and small intestine for analysis (Fig 3a). We found that CD4+ T-regulatory cell (Treg) numbers were increased in small intestinal lamina propria and not in the large intestine with administration of whole sauerkraut brine (Fig 3a). Treg numbers were similarly increased with intake of whole sauerkraut brine or filter-sterilized sauerkraut brine, indicating that fermented food metabolites alone have the capacity to promote regulatory responses in the small intestine (Fig 3b). We then administered lactate, an abundant fermented food metabolite, and found an increase in Treg as well (Fig 3c). Acetate, another abundant fermented food metabolite, did not lead to expanded Treg, indicating a distinct capacity of lactate to impact small intestinal immunity (Fig 3d).

**Figure 3:**
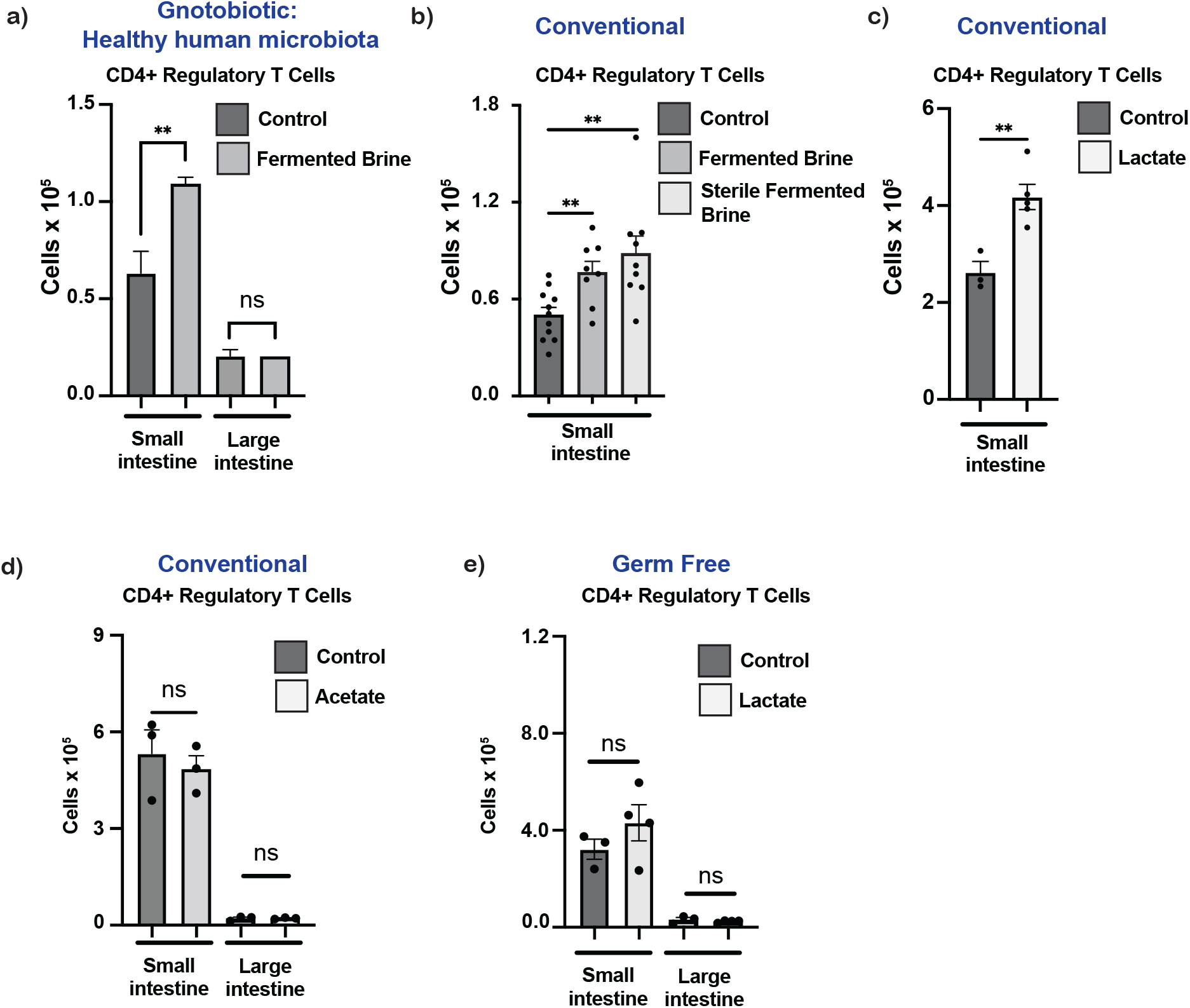
Metabolites of fermented foods promote microbiota dependent regulatory T cells. a)Gnotobiotic mice colonized with healthy human stool given sauerkraut brine for 4 weeks and CD4+Foxp3+ T regulatory cells isolated from the intestine were quantified b) Conventional mice given indicated treatment for 4-5 weeks and CD4+Foxp3+ T regulatory cells isolated from the intestine were quantified; results pooled across 3 experiments c) Conventional mice given lactate for 4 weeks and CD4+Foxp3+ T regulatory cells isolated from the intestine were quantified d) Conventional mice given acetate for 4 weeks and CD4+Foxp3+ T regulatory cells isolated from the intestine were quantified e) Germ Free mice given lactate for 4 weeks and T regulatory cells isolated from the intestine were quantified ** p ≤ 0.01, not significant (NS).

The regulatory T cell compartment of the intestine is regulated by the microbiota^17^ and we tested if the observed expansion of this cell type was dependent on the presence of a microbiome (Fig 3e). Indeed, lactate given to germ free mice did not significantly increase Treg numbers (Fig 3e). We then mono-colonized mice with a single fermented food-derived bacterial isolate with colonization of the small intestine (Fig 4a) and abundant lactate production, *Lacticaseibacillus paracasei*, to find that a single bacterial strain from fermented foods was able to confer increased regulatory T cells (Fig 4a). To clarify the role of bacterial metabolites in promoting regulatory T cells in a reductionist system, we mono-colonized mice with a single bacterial isolate with colonization of the small intestine (Fig 4b) and limited capacity to produce lactate, *E. coli* MG1655, and administered filter sterilized sauerkraut brine. E coli alone has no effect on Tregs in the small intestine (Fig 4b). However, when fermented food metabolites are administered, *E. coli* induced Treg cell response in the intestine is significantly increased (Fig 4b).

**Figure 4:**
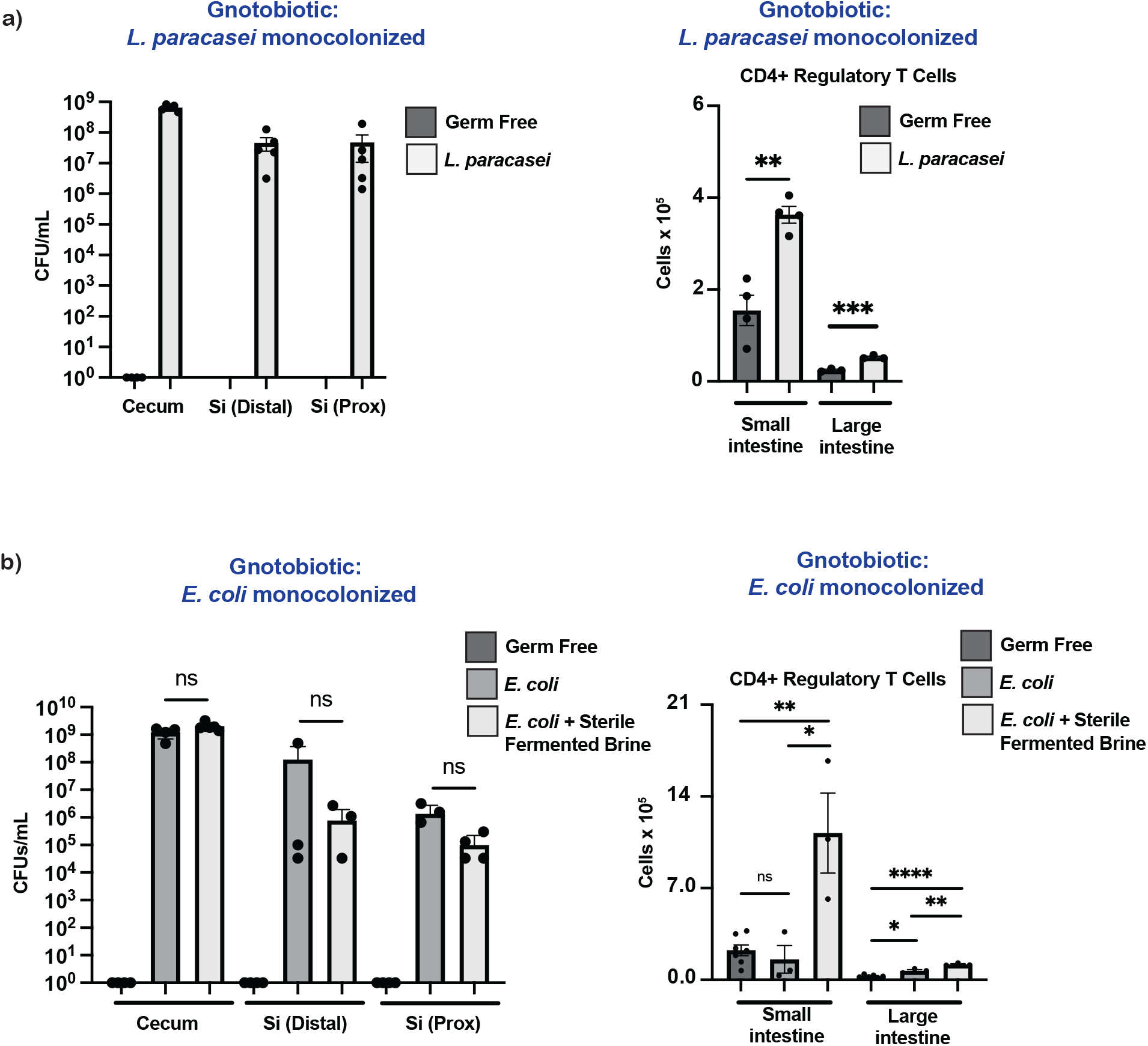
Either probiotic lactic acid bacteria or chemical metabolites from fermented foods promote regulatory T cell responses. a)Gnotobiotic mice colonized with a strain of fermented-food derived *L. paracasei* for 2 weeks. Left panel: Colony Forming Units (CFU) assessed in the site indicated, small intestine (Si), proximal (Prox). Right panel: CD4+, Foxp3+ T regulatory cells isolated from the intestine were quantified. b) Gnotobiotic mice colonized with *E. coli* MG1655 and treated with sterile fermented brine for 2 weeks. Left panel: Colony Forming Units (CFU) assessed in the site indicated, small intestine (Si), Proximal (Prox). Right panel: CD4+, Foxp3+ T regulatory cells isolated from the intestine were quantified. Not significant (NS), * p ≤ 0.05, ** p ≤ 0.01, *** <0.001, **** <0.0001.

## Discussion

Given the widespread and longstanding consumption of fermented foods paired with recent data suggesting they may confer health benefit, there is a pressing need to further characterize the microbial and chemical diversity as well as their effects on microbiome-host interactions^6^. The chemical diversity of fermented foods via an LC/MS microbial-metabolite focused pipeline, reveals grouping by fermentation type. The dominate organic acid metabolites that are the end products of fermentation, lactate and acetate, are inversely proportional, and their relative abundance corresponds to the broader metabolomic profile groups.

Recent findings demonstrate that fermented food intake alters gastrointestinal microbiota composition, however it remains unclear how live microbes or chemical metabolites independently interact with the microbiome^5–7^. We found that sterile sauerkraut brine or lactate alone altered the fecal and cecal microbiota of conventional mice in a distinct manner to whole brine, suggesting discrete roles of the components of fermented foods, specifically metabolites versus live microbial content, can impact gut microbes. When whole brine or lactate was given to germ free mice humanized with healthy human fecal sample, we observed a significant shift in the microbiota within the terminal ileum that was not apparent in feces, with enrichment of *Akkermanisa muciniphila*. Thus, in humanized mice, we see that the major impact of fermented food intake is exerted on the small intestinal microbiota and largely independent of live microbes.

Given fermented food promotion of *A. muciniphila* and the previously reported ability of this species to induce Tregs^18^, we investigated the host immune impact of fermented food. Notably, Tregs expand in the small intestinal lamina propria, but not in the large intestine. The site-specific alteration of microbiota composition and regulatory immune shift is consistent with oral intake and relatively higher concentrations of fermented food metabolites in the small intestine. Small intestinal Tregs were similarly expanded with delivery of sterile sauerkraut brine or lactate alone, but not with acetate. As Tregs in the intestine are regulated by the presence of the microbiome, we tested if the observed effect of lactate was microbiome-dependent. Indeed, germ-free mice that received lactate did not increase intestinal Tregs, which suggests that lactate is affecting microbiota dependent Tregs.

Colonization of germ free mice with a single bacteria known to colonize the small intestine and produce lactate, *Lacticaseibacillus paracasei*, increased Tregs in both the small and large intestine, likely because of increased lactate concentrations in the small and large intestine in this setting. Monocolonization of germ free mice with *E coli*, which colonizes the small intestine without significant lactate production, did not increase Tregs in the small intestine; however sterile fermented food given to *E coli*-colonized mice greatly expanded Tregs. These findings in a monoassociated gnotobiotic mouse model demonstrate that fermented foods can function as a diet derived signal to shift the tone of microbiota induced immunity independent of altered microbiota composition.

While we demonstrate that orally ingested lactate has potent host directed effects in the small intestine, the mechanism of this interaction remains unclear. We hypothesize that lactate will have direct host interaction, particularly in the small intestine. Lactate is used as an energy source by immune cells via the transport channels MCT1-4 and can also signal via GPR81 and GPR31^19, 20^. The promotion of Tregs by lactate in the tumor microenvironment is well studied and thought to involve both direct use of lactate as a carbon source by Tregs as a signaling capacity within myeloid cells ^21, 22, 23, 24^. The role of lactate in intestinal homeostasis is less well studied, however, recent data suggests that lactate signaling via GPR81 on hematopoietic cells is a critical regulator of inflammation^25^. Our results support that lactate is a keystone metabolite that contributes to intestinal immune regulation and that oral intake of lactate containing fermented foods may be a potent signal to limit inflammation in the small intestine via both microbiota and host direct effects.

Acetate, the fermentative end product of acetogenic fermentation in Kombucha, does not alter Tregs in the small intestine when orally ingested. This suggests that the critical regulatory function of acetate may be specific to its generation and sensing in the colon and is not augmented with oral intake^26–28^. While we did not assess the effects of whole or sterile Kombucha, our data demonstrate that acetate component of Kombucha does not likely promote immune regulation in the small intestine. It is likely that other fermented food metabolites within Kombucha do have host or microbiome effects, but this remains untested.

This study highlights the complexity and diversity among common fermented foods. While it has remained unclear the relative contribution of live microbes or microbial metabolites to the host effects of fermented foods, we find that microbial metabolites alone have the capacity to alter both microbiota composition and host immunity. Further, we find that fermented foods can be functionally segregated based on their dominant fermentative end product, lactate or acetate, with lactate containing fermented foods having potent immune regulatory capacity in the small intestine.

Similar to recent proposals for a daily intake of microbes, our findings suggest we may also benefit from a daily intake of bacterial metabolites and, in particular, lactate^29^. Future studies should focus on identifying specific metabolites within fermented foods that confer health benefits as well as the microbes and pathways that lead to their production^30^. Microbially and chemically defined fermented foods are poised to become powerful and delicious tools to promote health as well as prevent and treat disease.

## Author contributions

SPS and JLS conceived of experiments and wrote the manuscript. ELS, RHG, and SKH participated in and helped analyze mouse experiments. RNC and MMC aided with stool microbiome analysis. HCW, MW, and EBC aided with metabolomics analysis.

## Acknowledgments

This work was funded by generous donations from the Center for Human Microbiome Research. SPS was supported by NIH T32 DK007056. MMC was supported by Stanford Graduate Smith Fellowship. JLS. is a Chan-Zuckerberg Biohub investigator and supported by National Institutes of Health grant DP1-AT009892 and R01-DK085025. Cell sorting/flow cytometry analysis for this project was done on instruments in the Stanford Shared FACS Facility using NIH S10 Shared Instrument Grants S10RR025518-01, S10RR027431-01, 1S10OD026831-01. We wish to thank Arianna Burke for technical support.

## Data and materials availability

The authors declare that the data supporting the findings of this study are available within the paper and its supplementary information files.

## Materials and Methods

### DNA isolation and 16S amplicon sequencing

DNA was extracted from indicated source using the MoBio PowerSoil kit according to manufacturer’s protocol and amplified at the V4 region of the 16S ribosomal RNA (rRNA) subunit gene and 250 nucleotides (nt) Illumina sequencing reads were generated. 16S rRNA gene amplicon sequencing data was demultiplexed using the QIIME pipeline version 1.8^32^. Amplicon sequence variants (ASVs) were identified with a learned sequencing error correction model (DADA2 method)^33^ using the dada2 package in R. ASVs were assigned taxonomy using the GreenGenes database (version 13.8).

### Fermented foods

Fermented foods analyzed were commercially available products purchased from grocery stores.

### Gas chromatography/mass spectrometry (GC/MS) analysis of SCFAs and Lactate

*For Yogurt and Miso*, ~100 mg of semi-solid food was pre-weighed into a 2-ml screw top tube (or 96 well plate) containing six 6 mm ceramic beads (Precellys^®^ CK28 Lysing Kit). 500 μL of an extraction solution (20 μL 10 mM n-crotonic acid in water as internal standard, 100 μL 6 N HCl, 380 μL ddH_2_O) and 500 μL diethyl ether were then added to each tube. *For Sauerkraut Brine and Kombucha*, 1 ml of liquid food was centrifuged at 13000 × g for 10 min at room temperature. 100 μl of supernatant was added into 400 μl of extraction solution (20 μL 10 mM n-crotonic acid in water as internal standard, 100 μL 6 N HCl, 280 μL ddH_2_O) and 500 μL diethyl ether in the beads tube. *For authentic standards*, 100 μl of SCFAs/lactate mix solution (ranging from 5000 μM to 0.5 μM, series of 2-fold dilution) was added into 400 μl of extraction solution (20 μL 10 mM n-crotonic acid in water as internal standard, 100 μL 6 N HCl, 280 μL ddH_2_O) and 500 μL diethyl ether in the beads tube.

All samples were homogenized by vigorous shaking using a QIAGEN Tissue Lyser II at 25/s for 10 min. The resulting homogenates were subjected to centrifugation at 18000 x g for 10 min. The organic layer was transferred to a new glass vial (29391-U, Supelco) for derivatization using the following procedure: 100 μl of diethyl ether extract was mixed with 10 μL MTBSTFA and incubated at room temperature for 2 h. 1 μL of the derivatized samples were analyzed using a 7890B GC System (Agilent Technologies) and 5973 Network Mass Selective Detector (Agilent Technologies). We used the following chromatography conditions for GC-MS: Column: HP-5MS, 30 m, 0.25 mm, 0.25 μm; Injection Mode: splitless; Temperature Program: 40 °C for 0.1 min; 40-70 °C at 5 °C/min, hold at 70 °C for 3.5 min; 70-160 °C at 20 °C/min; 160-325 °C at 35 °C/min, equilibration for 3 min. 1 μL of each sample was injected and analyte concentrations were quantified by comparing their peak areas with those of external authentic standards using MassHunter Quantitative Analysis Software.

### Liquid chromatography-mass spectrometry (LC-MS)

Fermented food samples were extracted in LC-MS grade methanol. Sample supernatants were then transferred, evaporated, and reconstituted in an internal standard mix (50% Methanol). Metabolite samples were analyzed on a LC-MS qTOF instrument using reverse phase C18 positive, C18 negative, and HILIC positive methods as described^16^. Compound annotation was carried out using the MSDIAL software^31^ and an authentic standard reference library^16^. To quantify metabolite levels, area under the curve for each annotated metabolite was normalized using the sum of internal standards in each sample.

### Cell isolation from intestine

Protocol was adapted from prior^34,35^. Small intestine was removed from mice, Peyer’s patches were removed and tissue was cut longitudinally and washed in PBS. The tissue was then cut into 1-2 cm sections and placed into a 125 ml flask containing 20ml of RPMI containing 2% FBS, 5mM EDTA and 5mM DTT (DL-Dithiotreitol99%, Sigma-Aldrich catalog #D0632) and incubated for 20 min on stirring plate. The contents of the flask were then strained through a fine-meshed kitchen strainer into a 500 ml beaker. The pieces of small intestine were transferred to a 50 ml conical tube containing 10 ml of serum free media with 5mM EDTA and shaken vigorously for 30 seconds and then strained through the same kitchen strainer into the same beaker three times. The small intestine pieces recovered from strainer were washed with 5ml PBS and transferred into a 50 ml beaker, finely minced with surgical scissors, placed in 7ml of RPMI + 1mg/ml Liberase + 0.5mg/ml DNAse, and incubated for 25 min at 37°C on stirring plate. At end of incubation 5mls of RPMI containing 3% FBS was added and each sample was passed through a 70μm and 40μm filter with assistance from the plunger of a 3ml syringe.

### Antibodies and Flow cytometry

Antibodies for flow cytometry were purchased as listed below. Identification and gating of dead cells in all experiments was with LIVE/DEAD fixable dead stain (ThermoFisher). All stains were done out with anti-CD16/32 blocking antibody (clone 93, Biolegend). For intracellular staining, cells the FoxP3/Transcription factor staining buffer set was used according to the manufacturer’s directions (ThermoFisher). (**Antibody**/Fluorophore/Clone/Source): **CD45**/PerCP/Cyanine5.5/30-F11/BioLegend, **CD45.2**/Brilliant Violet 711/A20/BioLegend, **CD4**/Brilliant Violet 650/L3T4/BioLegend, **TCRb**/Alexa Fluor 700/H57-597/BioLegend, **TCRb**/PE-Cyanine7/H57-597/BioLegend, **CD8a**/Brilliant Violet 785/53-6.7/BioLegend, **T-bet**/PE-Cyanine7/4B10/Thermo Fisher, **Foxp3**/eFluor 450/FJK-16s/Thermo Fisher, **ROR gamma (t)**/PE/B2D/Thermo Fisher, **GATA3**/Alexa Fluor 488/L50-823/Thermo Fisher, **CD90.2**/BUV395/53-2.1/Thermo Fisher, **Ki67**/BD Horizon BV711/B56/Thermo Fisher, **IgA**/APC/mA-6E1/Thermo Fisher

### Mice

Conventional B6 mice were purchased from either Taconic Biosciences (B6NTac-EF) or The Jackson Laboratory. Mice were used between 6-12 weeks of age. All mice were bred and maintained under pathogen-free conditions at an American Association for the Accreditation of Laboratory Animal Care accredited animal facility. Fermented foods were administered either by daily gavage of 200ul for 4 weeks or delivered in the drinking water bottles of mice for 2-6 weeks duration.

### Statistical analysis

A two-tailed Student’s t-test was used for all statistical analysis unless otherwise noted, * p≤ 0.05, **p ≤ 0.01, *** p≤0.001, **** p≤0.0001, not significant (ns, p>0.05)

## References

1. Dimidi E, Cox SR, Rossi M, Whelan K. Fermented Foods: Definitions and Characteristics, Impact on the Gut Microbiota and Effects on Gastrointestinal Health and Disease. Nutrients. 2019;11(8). Epub 20190805. doi: 10.3390/nu11081806. PubMed PMID: 31387262; PMCID: PMC6723656.

2. Carrigan MA, Uryasev O, Frye CB, Eckman BL, Myers CR, Hurley TD, Benner SA. Hominids adapted to metabolize ethanol long before human-directed fermentation. Proc Natl Acad Sci U S A. 2015;112(2):458–63. Epub 20141201. doi: 10.1073/pnas.1404167111. PubMed PMID: 25453080; PMCID: PMC4299227.

3. Marco ML, Sanders ME, Gänzle M, Arrieta MC, Cotter PD, De Vuyst L, Hill C, Holzapfel W, Lebeer S, Merenstein D, Reid G, Wolfe BE, Hutkins R. The International Scientific Association for Probiotics and Prebiotics (ISAPP) consensus statement on fermented foods. Nat Rev Gastroenterol Hepatol. 2021;18(3):196–208. Epub 20210104. doi: 10.1038/s41575-020-00390-5. PubMed PMID: 33398112; PMCID: PMC7925329.

4. Lavefve L, Marasini D, Carbonero F. Microbial Ecology of Fermented Vegetables and Non-Alcoholic Drinks and Current Knowledge on Their Impact on Human Health. Adv Food Nutr Res. 2019;87:147–85. Epub 20181207. doi: 10.1016/bs.afnr.2018.09.001. PubMed PMID: 30678814.

5. Marco ML, Heeney D, Binda S, Cifelli CJ, Cotter PD, Foligné B, Gänzle M, Kort R, Pasin G, Pihlanto A, Smid EJ, Hutkins R. Health benefits of fermented foods: microbiota and beyond. Curr Opin Biotechnol. 2017;44:94–102. Epub 2016/12/18. doi: 10.1016/j.copbio.2016.11.010. PubMed PMID: 27998788.

6. Wastyk HC, Fragiadakis GK, Perelman D, Dahan D, Merrill BD, Yu FB, Topf M, Gonzalez CG, Van Treuren W, Han S, Robinson JL, Elias JE, Sonnenburg ED, Gardner CD, Sonnenburg JL. Gut-microbiota-targeted diets modulate human immune status. Cell. 2021;184(16):4137–53.e14. Epub 20210712. doi: 10.1016/j.cell.2021.06.019. PubMed PMID: 34256014.

7. Taylor BC, Lejzerowicz F, Poirel M, Shaffer JP, Jiang L, Aksenov A, Litwin N, Humphrey G, Martino C, Miller-Montgomery S, Dorrestein PC, Veiga P, Song SJ, McDonald D, Derrien M, Knight R. Consumption of Fermented Foods Is Associated with Systematic Differences in the Gut Microbiome and Metabolome. mSystems. 2020;5(2). Epub 20200317. doi: 10.1128/mSystems.00901-19. PubMed PMID: 32184365; PMCID: PMC7380580.

8. Tamang JP, Watanabe K, Holzapfel WH. Review: Diversity of Microorganisms in Global Fermented Foods and Beverages. Front Microbiol. 2016;7:377. Epub 20160324. doi: 10.3389/fmicb.2016.00377. PubMed PMID: 27047484; PMCID: PMC4805592.

9. Obafemi YD, Oranusi SU, Ajanaku KO, Akinduti PA, Leech J, Cotter PD. African fermented foods: overview, emerging benefits, and novel approaches to microbiome profiling. NPJ Sci Food. 2022;6(1):15. Epub 20220218. doi: 10.1038/s41538-022-00130-w. PubMed PMID: 35181677; PMCID: PMC8857253.

10. Leech J, Cabrera-Rubio R, Walsh AM, Macori G, Walsh CJ, Barton W, Finnegan L, Crispie F, O’Sullivan O, Claesson MJ, Cotter PD. Fermented-Food Metagenomics Reveals Substrate-Associated Differences in Taxonomy and Health-Associated and Antibiotic Resistance Determinants. mSystems. 2020;5(6). Epub 20201110. doi: 10.1128/mSystems.00522-20. PubMed PMID: 33172966; PMCID: PMC7657593.

11. Villarreal-Soto SA, Bouajila J, Pace M, Leech J, Cotter PD, Souchard JP, Taillandier P, Beaufort S. Metabolome-microbiome signatures in the fermented beverage, Kombucha. Int J Food Microbiol. 2020;333:108778. Epub 20200709. doi: 10.1016/j.ijfoodmicro.2020.108778. PubMed PMID: 32731153.

12. Macori G, Cotter PD. Novel insights into the microbiology of fermented dairy foods. Curr Opin Biotechnol. 2018;49:172–8. Epub 20170929. doi: 10.1016/j.copbio.2017.09.002. PubMed PMID: 28964915.

13. Landis EA, Oliverio AM, McKenney EA, Nichols LM, Kfoury N, Biango-Daniels M, Shell LK, Madden AA, Shapiro L, Sakunala S, Drake K, Robbat A, Booker M, Dunn RR, Fierer N, Wolfe BE. The diversity and function of sourdough starter microbiomes. Elife. 2021;10. Epub 20210126. doi: 10.7554/eLife.61644. PubMed PMID: 33496265; PMCID: PMC7837699.

14. Wolfe BE, Button JE, Santarelli M, Dutton RJ. Cheese rind communities provide tractable systems for in situ and in vitro studies of microbial diversity. Cell. 2014;158(2):422–33. doi: 10.1016/j.cell.2014.05.041. PubMed PMID: 25036636; PMCID: PMC4222527.

15. van de Wouw M, Walsh CJ, Vigano GMD, Lyte JM, Boehme M, Gual-Grau A, Crispie F, Walsh AM, Clarke G, Dinan TG, Cotter PD, Cryan JF. Kefir ameliorates specific microbiota-gut-brain axis impairments in a mouse model relevant to autism spectrum disorder. Brain Behav Immun. 2021;97:119–34. Epub 20210709. doi: 10.1016/j.bbi.2021.07.004. PubMed PMID: 34252569.

16. Han S, Van Treuren W, Fischer CR, Merrill BD, DeFelice BC, Sanchez JM, Higginbottom SK, Guthrie L, Fall LA, Dodd D, Fischbach MA, Sonnenburg JL. A metabolomics pipeline for the mechanistic interrogation of the gut microbiome. Nature. 2021;595(7867):415–20. Epub 20210714. doi: 10.1038/s41586-021-03707-9. PubMed PMID: 34262212.

17. Whibley N, Tucci A, Powrie F. Regulatory T cell adaptation in the intestine and skin. Nat Immunol. 2019;20(4):386–96. Epub 2019/03/19. doi: 10.1038/s41590-019-0351-z. PubMed PMID: 30890797.

18. Ansaldo E, Slayden LC, Ching KL, Koch MA, Wolf NK, Plichta DR, Brown EM, Graham DB, Xavier RJ, Moon JJ, Barton GM. induces intestinal adaptive immune responses during homeostasis. Science. 2019;364(6446):1179–84. doi: 10.1126/science.aaw7479. PubMed PMID: 31221858; PMCID: PMC6645389.

19. Llibre A, Grudzinska FS, O’Shea MK, Duffy D, Thickett DR, Mauro C, Scott A. Lactate cross-talk in hostpathogen interactions. Biochem J. 2021;478(17):3157–78. doi: 10.1042/BCJ20210263. PubMed PMID: 34492096; PMCID: PMC8454702.

20. Morita N, Umemoto E, Fujita S, Hayashi A, Kikuta J, Kimura I, Haneda T, Imai T, Inoue A, Mimuro H, Maeda Y, Kayama H, Okumura R, Aoki J, Okada N, Kida T, Ishii M, Nabeshima R, Takeda K. GPR31-dependent dendrite protrusion of intestinal CX3CR1. Nature. 2019;566(7742):110–4. Epub 20190123. doi: 10.1038/s41586-019-0884-1. PubMed PMID: 30675063.

21. Multhoff G, Vaupel P. Lactate-avid regulatory T cells: metabolic plasticity controls immunosuppression in tumour microenvironment. Signal Transduct Target Ther. 2021;6(1):171. Epub 20210430. doi: 10.1038/s41392-021-00598-0. PubMed PMID: 33931598; PMCID: PMC8087677.

22. Watson MJ, Vignali PDA, Mullett SJ, Overacre-Delgoffe AE, Peralta RM, Grebinoski S, Menk AV, Rittenhouse NL, DePeaux K, Whetstone RD, Vignali DAA, Hand TW, Poholek AC, Morrison BM, Rothstein JD, Wendell SG, Delgoffe GM. Metabolic support of tumour-infiltrating regulatory T cells by lactic acid. Nature. 2021;591(7851):645–51. Epub 20210215. doi: 10.1038/s41586-020-03045-2. PubMed PMID: 33589820; PMCID: PMC7990682.

23. Brown TP, Bhattacharjee P, Ramachandran S, Sivaprakasam S, Ristic B, Sikder MOF, Ganapathy V. The lactate receptor GPR81 promotes breast cancer growth via a paracrine mechanism involving antigen-presenting cells in the tumor microenvironment. Oncogene. 2020;39(16):3292–304. Epub 20200219. doi: 10.1038/s41388-020-1216-5. PubMed PMID: 32071396.

24. Colegio OR, Chu NQ, Szabo AL, Chu T, Rhebergen AM, Jairam V, Cyrus N, Brokowski CE, Eisenbarth SC, Phillips GM, Cline GW, Phillips AJ, Medzhitov R. Functional polarization of tumour-associated macrophages by tumour-derived lactic acid. Nature. 2014;513(7519):559–63. Epub 20140713. doi: 10.1038/nature13490. PubMed PMID: 25043024; PMCID: PMC4301845.

25. Ranganathan P, Shanmugam A, Swafford D, Suryawanshi A, Bhattacharjee P, Hussein MS, Koni PA, Prasad PD, Kurago ZB, Thangaraju M, Ganapathy V, Manicassamy S. GPR81, a Cell-Surface Receptor for Lactate, Regulates Intestinal Homeostasis and Protects Mice from Experimental Colitis. J Immunol. 2018;200(5):1781–9. Epub 20180131. doi: 10.4049/jimmunol.1700604. PubMed PMID: 29386257; PMCID: PMC5858928.

26. Takeuchi T, Miyauchi E, Kanaya T, Kato T, Nakanishi Y, Watanabe T, Kitami T, Taida T, Sasaki T, Negishi H, Shimamoto S, Matsuyama A, Kimura I, Williams IR, Ohara O, Ohno H. Acetate differentially regulates IgA reactivity to commensal bacteria. Nature. 2021;595(7868):560–4. Epub 20210714. doi: 10.1038/s41586-021-03727-5. PubMed PMID: 34262176.

27. Smith PM, Howitt MR, Panikov N, Michaud M, Gallini CA, Bohlooly YM, Glickman JN, Garrett WS. The microbial metabolites, short-chain fatty acids, regulate colonic Treg cell homeostasis. Science. 2013;341(6145):569–73. doi: 10.1126/science.1241165. PubMed PMID: 23828891; PMCID: 3807819.

28. Maslowski KM, Vieira AT, Ng A, Kranich J, Sierro F, Yu D, Schilter HC, Rolph MS, Mackay F, Artis D, Xavier RJ, Teixeira MM, Mackay CR. Regulation of inflammatory responses by gut microbiota and chemoattractant receptor GPR43. Nature. 2009;461(7268):1282–6. doi: 10.1038/nature08530. PubMed PMID: 19865172; PMCID: PMC3256734.

29. Marco ML, Hill C, Hutkins R, Slavin J, Tancredi DJ, Merenstein D, Sanders ME. Should There Be a Recommended Daily Intake of Microbes? J Nutr. 2020;150(12):3061–7. doi: 10.1093/jn/nxaa323. PubMed PMID: 33269394; PMCID: PMC7726123.

30. Marco ML. Defining how microorganisms benefit human health. Microb Biotechnol. 2021;14(1):35–40. Epub 20201025. doi: 10.1111/1751-7915.13685. PubMed PMID: 33099885; PMCID: PMC7888441.

31. Tsugawa H, Cajka T, Kind T, Ma Y, Higgins B, Ikeda K, Kanazawa M, VanderGheynst J, Fiehn O, Arita M. MS-DIAL: data-independent MS/MS deconvolution for comprehensive metabolome analysis. Nat Methods. 2015;12(6):523–6. Epub 20150504. doi: 10.1038/nmeth.3393. PubMed PMID: 25938372; PMCID: PMC4449330.

32. Caporaso JG, Kuczynski J, Stombaugh J, Bittinger K, Bushman FD, Costello EK, Fierer N, Peña AG, Goodrich JK, Gordon JI, Huttley GA, Kelley ST, Knights D, Koenig JE, Ley RE, Lozupone CA, McDonald D, Muegge BD, Pirrung M, Reeder J, Sevinsky JR, Turnbaugh PJ, Walters WA, Widmann J, Yatsunenko T, Zaneveld J, Knight R. QIIME allows analysis of high-throughput community sequencing data. Nat Methods. 2010;7(5):335–6. Epub 2010/04/11. doi: 10.1038/nmeth.f.303. PubMed PMID: 20383131; PMCID: PMC3156573.

33. Callahan BJ, McMurdie PJ, Rosen MJ, Han AW, Johnson AJ, Holmes SP. DADA2: High-resolution sample inference from Illumina amplicon data. Nat Methods. 2016;13(7):581–3. doi: 10.1038/nmeth.3869. PubMed PMID: 27214047; PMCID: 4927377.

34. Goodyear AW, Kumar A, Dow S, Ryan EP. Optimization of murine small intestine leukocyte isolation for global immune phenotype analysis. J Immunol Methods. 2014;405:97–108. Epub 20140204. doi: 10.1016/j.jim.2014.01.014. PubMed PMID: 24508527.

35. Spencer SP, Wilhelm C, Yang Q, Hall JA, Bouladoux N, Boyd A, Nutman TB, Urban JF, Jr., Wang J, Ramalingam TR, Bhandoola A, Wynn TA, Belkaid Y. Adaptation of innate lymphoid cells to a micronutrient deficiency promotes type 2 barrier immunity. Science. 2014;343(6169):432–7. Epub 2014/01/25. doi: 10.1126/science.1247606. PubMed PMID: 24458645.

